# Experimental and mathematical insights on the competition between poliovirus and a defective interfering genome

**DOI:** 10.1101/519751

**Authors:** Yuta Shirogane, Elsa Rousseau, Jakub Voznica, Igor M. Rouzine, Simone Bianco, Raul Andino

**Affiliations:** Department of Microbiology and Immunology, University of California, San Francisco, San Francisco, CA 94158, USA; Department of Virology, Faculty of Medicine, Kyushu University, Fukuoka, 812-8582, Japan; Department of Industrial and Applied Genomics, S2S - Science to Solution, IBM Almaden Research Center, 650 Harry Road, San Jose, CA 95120-6099, USA; NSF Center for Cellular Construction, University of California, San Francisco, San Francisco, CA 94158, USA; ENS Cachan, Université Paris-Saclay, 61 Avenue du Président Wilson, Cachan, 94230, France; UMR 7238 CNRS, Université Pierre et Marie Curie, Institut de Biologie Paris Seine, France

## Abstract

During replication, RNA virus populations accumulate genome alterations, such as mutations and deletions. The interactions between individual variants within the population can determine the fitness of the virus and, thus, the outcome of infection. We developed an ordinary differential equation model to infer the effect of the interaction between defective interfering (DI) replicons and wild-type (WT) poliovirus. We measure production of RNA and viral particles during a single infection cycle, and use these data to infer model parameters. We find that DI replicates faster than WT, but an equilibrium is established when both WT and DI compete for resources needed for RNA replication and genome encapsidation. In the presence of DI, the concentration of WT virions at cell lysis is suppressed by the factor of 5. Multiple generations within a single cell infection provide opportunities for significant inhibition of WT replication by competition with the faster replicating DI genomes.

## Introduction

Co-infections, the simultaneous infection of a host by multiple pathogen species, are frequently observed (***Mideo, 2009; Read and Taylor, 2001***). The interactions between these microrganisms can determine the trajectories and outcomes of infection. Indeed, competition between pathogen species or strains is a major force driving the composition, dynamics and evolution of such populations (***Mideo, 2009; Bashey, 2015***). Three types of competition among free-living organisms have been defined from an ecological point of view: exploitation, apparent and interference competition (***Read and Taylor, 2001; Bashey, 2015; Mideo, 2009***). Exploitation competition is a passive process in which pathogens compete for access to host resources. Apparent competition is competition that is not due to using shared resources, but to having a predator in common (***Holt, 1977***), and is generally associated with the stimulation of host immune response (***Bashey, 2015***). Interference competition represents a direct attack inhibiting the growth, reproduction or transmission of competitors, either chemically or mechanically (***Schoener, 1983***).

Here we focus on the interference competition between two RNA virus genomes co-infecting a single cell, one being a fully operational wild-type (WT) poliovirus type 1 (PV1) and the other a defective replicon of PV1, lacking the region encoding for capsid proteins, and called a defective interfering (DI) genome. Poliovirus, the causative agent of poliomyelitis, is a positive-sense singlestranded RNA enterovirus belonging of the family *Picornaviridae* (***Racaniello, 2013***). Upon cell infection, WT poliovirus initiates a series of processes that leads to the production of structural and nonstructural viral proteins and genome amplification. Structural proteins encapsidate the viral RNA, which leads to infectious viral particle production and cell-to-cell spread. Indeed, only encapsidated poliovirus genomes can survive outside the cells and can bind to new cells to initiate infections.

As an RNA virus, poliovirus is characterized by a high level of genome plasticity and evolution capacity, due to both high replication rate and error-prone nature of viral RNA polymerase (***Holland et al., 1982; Drake, 1993***), which generates a large proportion of mutants in the viral population, called viral quasispecies (***Nowak, 1992; Eigen, 1993; Biebricher and Eigen, 2006***). In addition, defective genomes, lacking essential genes, are produced by defective replication or viral RNA recombination (***von Magnus, 1954; Hirst, 1962; Sergiescu et al., 1966***). Natural poliovirus DI particles have been observed with in-frame deletion in the P1 region of the genome encoding structural capsid proteins, while expression of nonstructural proteins is not affected (***Kuge et al., 1986; Lundquist et al., 1979***). The DI genome used in this study features similar deletion of the P1 region. When co-infecting a cell with the WT, it can exploit capsid proteins produced by the WT to form DI particles, and in this way spread to new cells. However, it is not able to reproduce when not co-infecting a cell with the WT helper virus (***Lundquist et al., 1979***).

From an ecological perspective, the WT can be viewed as a cooperator, producing capsid proteins as public goods. The DI particles are non-producing cheaters, bearing no production cost while exploiting the capsid protein products from the WT (***Mideo, 2009; Bashey, 2015***). Hence, co-infection should enable DI particles to replicate and spread, while hindering WT growth and propagation by interference competition.

DI particles have been reported for a large number of viral species, such as vesicular stomatitis virus (***Bellett and Cooper, 1959***), poliovirus (***Cole et al., 1971***), ebola virus (***Calain et al., 1999***), dengue virus (***Li et al., 2011***) or influenza virus (***Saira et al., 2013; Huang, 1973***). Given that DI particles hamper WT production, they can attenuate WT virus infection (***Huang, 1973; Cole et al., 1971***). A recent study showed that defective viral genomes can contribute to attenuation in influenza virus infected patients (***Vasilijevic et al., 2017***), and they can protect from experimental challenges with a number of pathogenic respiratory viruses (***Dimmock and Easton, 2015; Sun et al., 2015***).

In this study, we examined the competition between WT and DI PV1 genomes within cells during one infection cycle. We then developed an ordinary differential equations (ODEs) mathematical model to capture the dynamics of DI and WT replication and encapsidation. We aimed to understand the mechanisms of interference of the defective genome with WT within a cell. Our study indicates that DI and WT genomes compete for limiting cellular resources required for their genome amplification and for capsid proteins required for particle morphogenesis. The model was further used to evaluate the potential outcome of the interaction between DIs and WT viruses over a large range of parameter values and initial conditions (multiplicities of infection, temporal spacing and order of infection). As a result we identified the most important parameters affecting WT production by DI co-infection. DI particles are spontaneously generated during a significant number of virus infection, but the factors affecting DIs generation and propagation are not well understood. Our results indicate that DIs compete a two different steps during poliovirus replication. This competition affects virus production and presumable the outcome of infection and pathogenesis. Thus, the mathematical model describe here will facilitate the understanding the biological significance of defective interference particles in the context of virus infection.

## Results

### Interference of WT poliovirus production by DI genomes

Initially, we evaluated whether DI genomes, carrying a deletion of the entire region encoding for capsid proteins, could affect progression of WT virus infection (***Figure 1***A). The DI genome used in this study does not produce capsid proteins and, thus, it is unable to encapsidate its genome and spread to other cells. However, it retains full capacity to produce non-capsid viral proteins and replicate its genomic RNA. WT poliovirus and DI genomic RNAs were transfected by electroporation to HeLaS3 cells and infectious titers of WT virus were determined over time by plaque assay (Material and Methods). As a control we also evaluated a replication-incompetent defective RNA lacking the capsid-encoding region, a part of 3D-polymerase encoding region, and the entire 3′ nontranslated region (NTR). HeLaS3 cells transfected by only WT genomes produced nearly 1×10^7^ PFU/ml WT virus 9 hours after transfection, while co-transfection of WT genomes together with DI RNAs resuted in 100-fold decrease of WT titers (***Figure 1***B). The non-replicating defective RNA did not affect WT virus production, suggesting that replicating DI genomes are required for effective interference, as previously reported (***Kaplan and Racaniello, 1988***).

### Quantification of the copy number of WT and DI genomes following co-transfection

Next, we examined the interaction between DI and WT genomes by varying the ratio of each RNA used to initiate transfection. Starting with equal RNA concentrations (5*µ*g) DI genomes were 4 times more efficiently transfected than WT (data not shown). Given that DI genomes are ∼2,000 nucleotide shorter (∼1/4 shorter) than WT genomes, the copy number of DI genomes are higher than that of WT genomes and transfection of shorter genomes is also more efficient than larger RNAs. We optimized our protocol to deliver equal copy number of DI and WT genomes into the transfected cells. We transfected 5*µ*g of WT to 1.25*µ*g of DI genomes, and we collected RNA samples at given timepoints (t=0, 2, 3.5, 5, 7 and 9 hours after transfection). The average number of genomes in a single cell was estimated as the number of genomes divided by the number of transfected cells. Replication rates of WT and DI decreased ∼7 hours after co-transfection, but this effect was not observed in the cells transfected only with WT or DI (***Figure 1***C). Thus, replication of WT genomes was inhibited by DI genomes, and the number of accumulated DI also decreased in the presence of WT. This suggests that WT and DI genomes compete for a limiting factor for replication. Nonetheless, DI genomes replicated faster than WT genomes (***Figure 1***C). To determine the numbers of encapsidated WT and DI genomes, we also treated cell lysates with a mixture of RNase A and RNase T1. Viral RNAs encapsidated in virus particles are resistant to RNase activity, while naked RNAs are degraded by RNase-treatment. The decrease of encapsidated WT genomes between singly and dually infected cells conditions was two-fold larger than that of WT genomes without RNase-treatment, indicating that DI genomes hamper WT genome encapsidation (***Figure 1***C&D, compare the difference between plain and dashed blue lines at 9 hours after transfection in ***Figure 1***D*i* to the difference in ***Figure 1***C*i*).

Thus, these results are consistent with the idea that DI RNAs replicate faster than WT genomes, due to their shorter genome (***Holland, 1991; Chao and Elena, 2017***). Interestingly, co-transfection results in a net reduction in replication of both WT and DI genomes most likely due to competition for some host-cell limiting factor needed for genome amplification. In addition, capsid proteins, produced by WT genomes, limit DI and WT virus production as DI genomes compete for these proteins and thus further inhibit WT production. To further examine the mechanism of defective interference and quantitatively evaluate the effect of co-replicating DIs, we designed a simple mathematical model that describes the DI/WT genome interactions.

## Mathematical description of intracellular competition

A deterministic mathematical model describing the intracellular competition between DI and WT genomes was developed, adapted from an existing competition model for human immun-odeficiency virus (HIV) (***Rouzine and Weinberger, 2013***). In order to describe appropriately the intracellular dynamics of poliovirus, we explicitly account for limiting resources depleted by the virus during replication, slowing down the growth of the population over time. The effect of limiting resources on poliovirus replication has been investigated by ***Regoes et al.*** (***2005***) using a mathematical model to explain the observed saturation in viral replication dynamics. Resources depleted by the virus may include phospholipids and *de novo* synthesized membranes for the formation of replication organelles (***Guinea and Carrasco, 1990; Nchoutmboube et al., 2013***), host factors regulating viral replication (***Altan-Bonnet, 2017***), or, as theoretically hypothesized, the number of ribosome complexes available for translation, the supply of amino acids for building proteins or the supply of nucleotides (***Regoes et al., 2005***). Our model considers a generic set of resources (*R*) at the virus disposal during replication. It describes the changes, over the course of infection of a cell, of the numbers of WT (*G*_*WT*_) and DI (*G*_*DI*_) positive-sense RNA genome copies, of free capsids produced by the WT (*C*) and of limiting resource units (*R*) depleted by the genomes for their replication. The set of ODEs is the following:

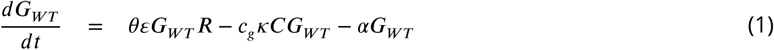

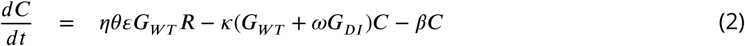

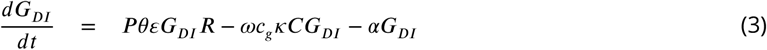

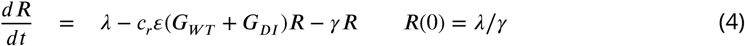

**Figure 1.**
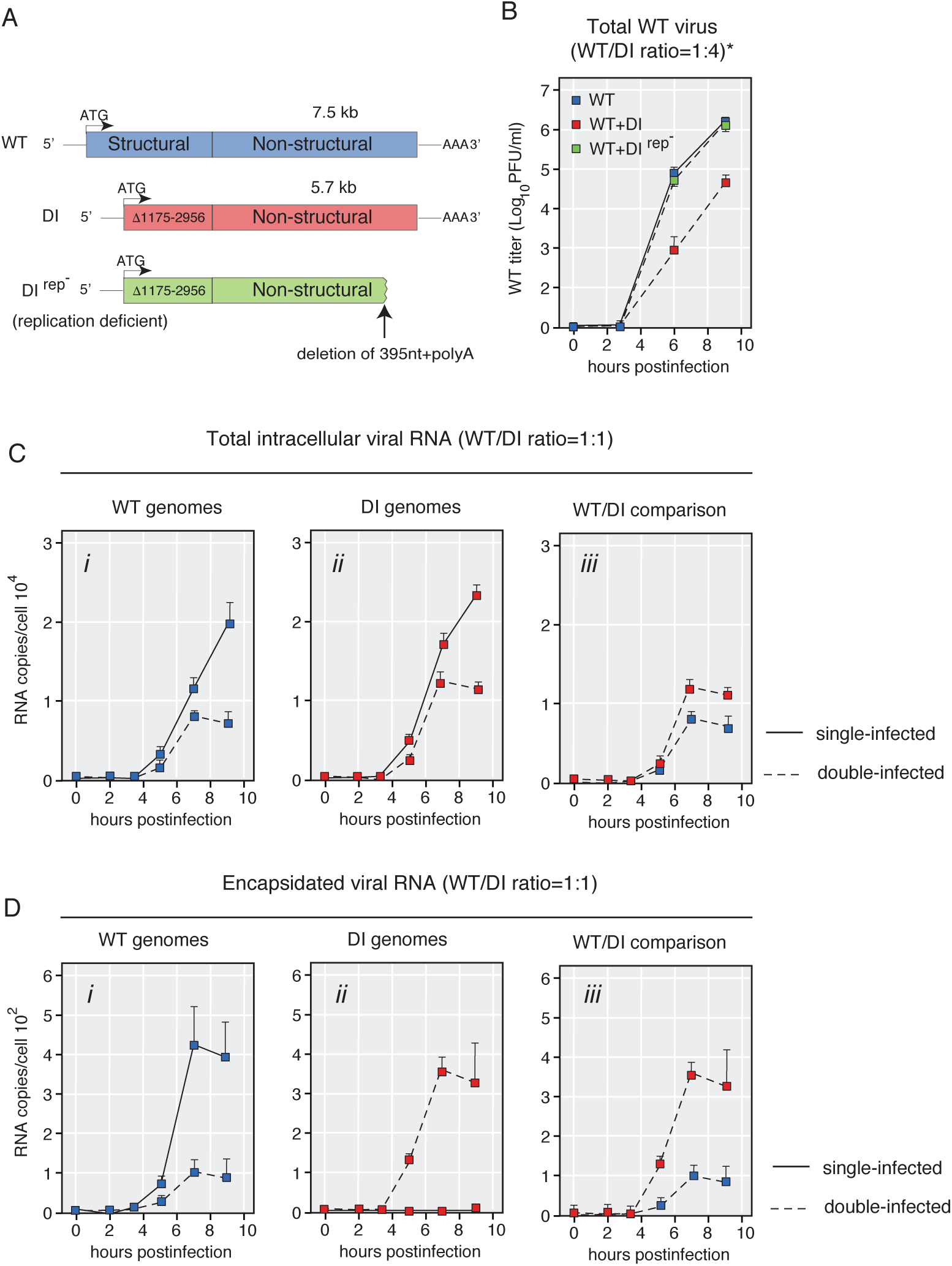
DI genomes inhibit WT virus production via genomic replication and encapsidation. (A) Structure of the WT poliovirus, DI(Δ1175-2956), and DI(Δ1175-2956)(Rep-). The DI genome has an in-frame deletion in its P1-encoding region. The PvuII-cut DI genome is used for a non-replicating RNA control (Rep-). (B) Growth curves of WT poliovirus after transfection of WT genomes and/or DI genomes. Genomic RNA was transfected to HeLaS3 cells, and samples were collected at t=0, 3, 6, and 9 hours after transfection. Virus titers were determined by plaque assay as described in Material and Methods. (C, D) Each or both of WT genomes and DI genomes were transfected to HeLaS3 cells. Copy numbers of total (C) or encapsidated (D) genomes were determined over time after transfection as described in Material and Methods. Blue and red squares indicate copy numbers of WT and DI genomes, respectively. Solid lines indicate single-infected (WT or DI genome-transfected) samples, while dotted lines indicate double-infected (both WT and DI genomes-transfected) samples.

Model parameters are summarized in ***Table 1*** and can be described through three distinct stages of the viral cycle: replication, capsid synthesis and encapsidation. A flow diagram of the model is available in ***Figure 2***. Limiting resources are produced at a linear rate *λ* and captured by WT (*G*_*WT*_) and DI (*G*_*DI*_) genomes at a rate *ε* per unit of resource (uor) per minute. One uor and one viral genome, by definition, form one replication complex (*c*_*r*_ = 1 uor • genome^-1^, ***den Boon and Ahlquist*** (***2010***)). Conditionally on the capture of a resource unit, a WT genome replicates and turn into *θ* genomes, before the replication complex disintegrates (we set the condition *θ* > 1 in order for virus genomes to replicate). We assume that this happens quickly compared to the other processes. DI genomes replicate faster than WT genomes by a fixed factor *P* (*P* > 1), which can be at least partially explained by the smaller genome size of the DI (***Holland, 1991; Chao and Elena, 2017***). The replication rate represents an average over the three main steps of poliovirus replication: (i) translation of the positive-sense RNA genome (***Novak and Kirkegaard, 1994***), (ii) transcription into negative-sense RNA genome that will be used as template for (iii) transcription into new positive-sense RNA genomes (***Semler and Wimmer, 2002; Regoes et al., 2005***). Because DI genomes lack the genes responsible for capsid proteins synthesis, only WT genomes are capable of producing free capsids (*C*), with the capsid-to-genome accumulation ratio *η*. WT genomes are then encapsidated (i.e. packaged into free capsids) at rate *κ*. DI genomes are assumed to encapsidate faster by a fixed factor *ω* (*ω* > 1). One viral genome encapsidates into one capsid to form a virion (*c*_*g*_ = 1 genome • capsid^−1^). Finally, *α, β*, and *γ* represent the decay rates of, respectively, viral genomes, free capsids and resources.

**Table 1.** Notations used in the model and model parameters.

| Notation | Definition | Unit<sup>a</sup> |  |  |  |
| --- | --- | --- | --- | --- | --- |
| **Observed variables** |  |  |  |  |  |
| $g_{WT}^{tot}, g_{DI}^{tot}$ | WT, DI total genome number | gen | | | |
| $v_{WT}, v_{DI}$ | WT, DI virion number | gen | | | |
| **State variables** |  |  |  |  |  |
| $G_{WT}, G_{DI}$ | WT, DI genome number | gen | | | |
| $C$ | Free capsid number | caps | | | |
| $R$ | Resource units | uor | | | |
| $C_{WT}, C_{DI}$ | WT, DI virion number | gen | | | |
| **Model parameters<sup>b</sup>** |  |  | **Best value<sup>c</sup>** | **Confidence Interval<sup>d</sup>** | **Sensitivity analysis<sup>e</sup>** |
| $c_g$ | Number of viral genomes per capsid (1) | gen · caps<sup>-1</sup> | | | |
| $c_r$ | Number of uor per viral genome (1) | uor · gen<sup>-1</sup> | | | |
| $\theta$ | Genome replication factor | dimensionless | 1: 7.297<br>2: 16.122 | [1.000; 28.87] | 1: [3, 5, 7, 9, 11]<br>2: [8, 12, 16, 20, 24] |
| $\epsilon$ | Resource capture rate by viral genomes | uor<sup>-1</sup> · min<sup>-1</sup> | $1.814 \cdot 10^{-6}$ | $[1.753; 1.877] \times 10^{-6}$ | $[0.8, 1.3, 1.8, 2.3, 2.8] \times 10^{-6}$ |
| $\kappa$ | Encapsidation rate | gen<sup>-1</sup> · min<sup>-1</sup> | $5.097 \cdot 10^{-6}$ | $[4.900; 5.282] \times 10^{-6}$ | $[2, 3.5, 5, 6.5, 8] \times 10^{-6}$ |
| $\alpha$ | Viral genome decay rate | min<sup>-1</sup> | 0 | - | 0 |
| $\eta$ | Capsid to genome accumulation ratio | caps · gen<sup>-1</sup> | $5.260 \cdot 10^{-2}$ | $[5.058; 5.439] \times 10^{-2}$ | $[2, 3.5, 5, 6.5, 8] \times 10^{-2}$ |
| $\beta$ | Capsid decay rate | min<sup>-1</sup> | $2.589 \cdot 10^{-3}$ | $[1.659; 3.476] \times 10^{-3}$ | $[1, 1.75, 2.5, 3.25, 4] \times 10^{-3}$ |
| $P$ | DI-to-WT replication ratio | dimensionless | 1.075 | [1.062; 1.087] | [1, 1.15, 1.3, 1.45, 1.6] |
| $\omega$ | DI-to-WT encapsidation ratio | dimensionless | 2.185 | [1.943; 2.453] | [1, 1.75, 2.5, 3.25, 4] |
| $\lambda$ | Resource linear production rate | uor · min<sup>-1</sup> | $1.000 \cdot 10^{-3}$ | $[1.000 \cdot 10^{-3}; 1.527 \cdot 10^{-1}]$ | $[1, 1.25, 1.5, 1.75, 2] \times 10^{-3}$ |
| $\gamma$ | Resource decay rate | min<sup>-1</sup> | 1: $4.381 \cdot 10^{-7}$<br>2: $9.680 \cdot 10^{-7}$ | $[1.583 \cdot 10^{-7}; 2.853 \cdot 10^{-5}]$ | 1: $[2, 3.25, 4.5, 5.75, 7] \times 10^{-7}$<br>2: $[4.5, 7, 9.5, 12, 14.5] \times 10^{-7}$ |
| $L$ | Logistic's maximum | dimensionless | $3.078 \cdot 10^{-2}$ | $[3.061; 3.099] \times 10^{-2}$ | - |
| $s$ | Logistic's steepness | dimensionless | $-3.234 \cdot 10^{-2}$ | $[-3.234; -3.234] \times 10^{-2}$ | - |
| $t_0$ | Logistic's midpoint | min | 318.459 | [318.459; 318.459] | - |

<sup>a</sup> gen: genome; caps: capsid; uor: unit of limiting resource<sup>b</sup> Fixed parameter values in parenthesis<sup>c</sup> Best optimized value shared by the 85 first sets (identical until fourth significant digits) for reduced model optimization ( $P, \omega, \kappa, \eta, \alpha, \beta, L, s, t_0$ ), and two arbitrarily chosen best sets for full model optimization ( $\epsilon, \theta, \lambda, \gamma$ ), the first set with $\theta$ value falling in the peak of its best values distribution (see **Figure 3–Figure Supplement 1A**).<sup>d</sup> Range of variation over 150 (reduced model) or 123 (full model) best sets of parameter values<sup>e</sup> For parameters $\theta$ and $\gamma$ , two vectors were tested, built on the two parameter estimations giving rise to the same high fit quality of the model to experimental data. As no noticeable differences were detected between both conditions, only those with small $\theta$ are presented in this paper.

**Figure 2.**
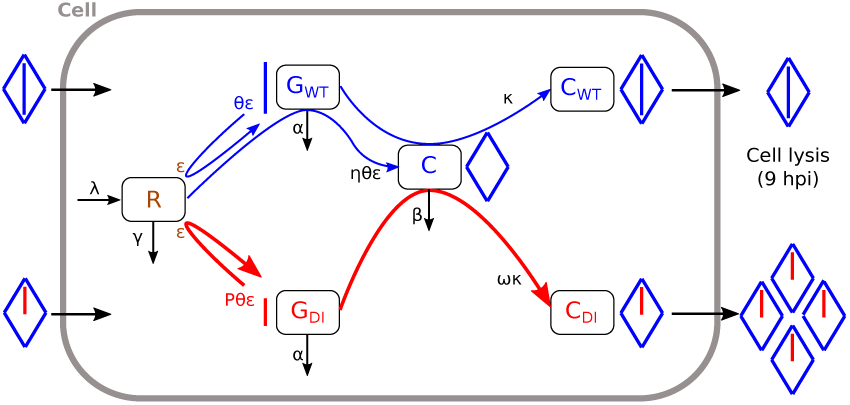
Flow diagram of the model (***Equation 1***–***Equation 6***).

The number of encapsidated WT genomes (i.e. WT virions, *C*_*WT*_) and DI genomes (i.e. DI virions, *C*_*DI*_) were measured experimentally (***Figure 1***D) and can easily be derived from ***Equation 2*** as the loss of free capsids due to encapsidation:

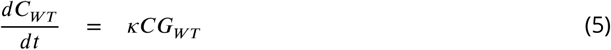

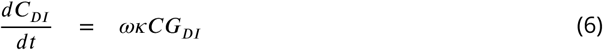

Further, burst sizes, which are defined as the number of virions at cell lysis, i.e. 9 hours post transfection (hpt, ***Burrill et al.*** (***2013***); ***Bird et al.*** (***2014***)), can be written as ℬ _*WT*_ = C_*WT*_(9 hpt) for WT virions and ℬ_*DI*_ = *C*_*DI*_ (9 hpt) for DI virions.

The system ***Equation 1***– ***Equation 6*** encompasses 10 parameters and six variables, among which only four variables could be experimentally measured. Therefore, the mathematical model presents a classical problem of parameter identifiability, specifically regarding parameters *ε* and *θ* that appear as a product, with parameter *ε* figuring separately in ***Equation 4*** corresponding to the unmeasured variable *R*. To solve this problem, we built a reduced model by assuming that the decrease in resources due to viral uptake follows a logistic decreasing function. Using a two-step optimization procedure, we are able to estimate the model parameters with reasonably high confidence (see Material and Methods and ***Table 1***).

Model predictions fit well the experimental measurements in ***Figure 1***, with best *R*^2^ = 0.974 for the reduced model and 0.965 for the full model (***Figure 3***). In dually infected cells, both versions of the model reproduce well the number of naked and encapsidated genome copies. In singly infected cells, the reduced model somewhat underestimates the number of naked genome copies while fitting satisfactorily the number of encapsidated genome copies. Conversely, the full model describes well the number of naked genome copies but overestimates the number of encapsidated genome copies. Overall, either model is able to capture the lower genome production when WT and DI coinfect a cell as compared to singly infected cells (compare same color plain curves between ***Figure 3***A & B). We hypothesized that this effect is the consequence of competing for limiting resources necessary for replication. The model also reproduces the fact that the WT is more hindered by this competition for resources than the DI (***Figure 3*** A, compare red and blue plain curves) thanks to the higher replication rate of DI genomes (*P* = 1.075, ***Table 1***). Predictions also demonstrate that DI genomes are more efficiently encapsidated than WT genomic RNA (***Figure 3***C, compare red and blue plain curves), thanks to the higher encapsidation rate of DI genomes (*ω* = 2.185, ***Table 1***). Most importantly, the model is able to describe the decrease in WT encapsidated genomes in dual infection compared to single WT infections (compare ***Figure 3***C & D, blue curves). Thus, it predicts a strong interference between DI and WT.

All parameter estimates are narrowly defined by the fitting procedure, except for *θ, λ* and *γ* (***Table 1***), as they tend to correlate with each other, yielding non-uniqueness of best-fit parameter values (***Figure 3***–***Figure Supplement 1***A,C-D, see also the distribution of *ε* values in ***Figure 3***–***Figure Supplement 1***B). Also, a strong log-to-log relationship was found between the resource production to decay ratio *λ* /*γ* and the replication factor *θ* (***Figure 3***–***Figure Supplement 1***E).

Both models feature a time-dependent virus replication rate (***Figure 3***–***Figure Supplement 1***F). In the reduced model, it is given by the logistic function Λ (*t*) (***Equation 10*** in Material and Methods), and in the full model by the product *θ ε R*(*t*). In both models, the best fit yields approximately the same time-dependent replication rate, starting at 3.07 • 10^−2^ for the reduced model and at 3.02 • 10^−2^ (*θ ε λ* /*γ*) for the full model, and decreasing with time towards 0. According to the reduced model, the time of half-decay is 318 minutes (*t*_0_ in ***Table 1***), which corresponds to 5.3 hours post transfection.

The predictive power of the full model with best estimated parameter values was tested on independent experimental measurements of WT burst size corresponding to various initial DIto-WT ratios for which the model had not been trained (***Figure 3***–***Figure Supplement 2***). Relative experimental and predicted WT burst sizes were normalized by their respective value for WT-only infection. The model was able to predict experimental outputs fairly well, albeit underestimating WT output for some DI-to-WT input ratios. The largest underestimation was observed for the DI-to-WT input ratio of 0.5, and this discrepancy vanished as the input ratio increased.

**Figure 3.**
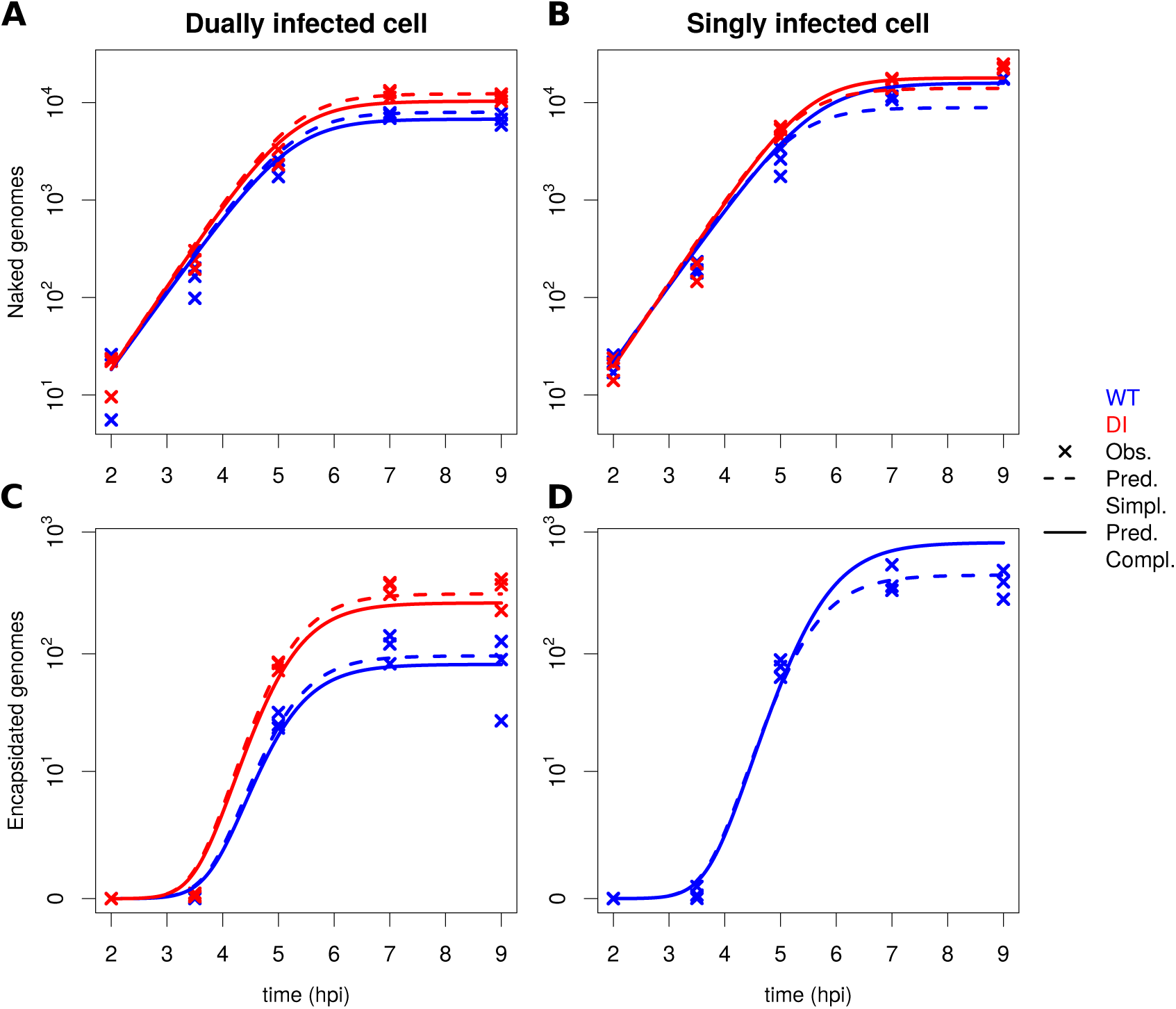
Experimental data and model predictions for the naked and encapsidated wild-type (WT) and defective interfering (DI) genomes. Evolution of the number of WT and DI (A-B) naked genome copies and (C-D) encapsidated genome copies with time, from 2 to 9 hours post transfection (hpt). (A & C) show data in dually infected cells whereas (B & D) show data in singly infected cells. WT and DI results are shown in blue and red color, respectively. Symbols x indicate experimental data for 3 replicates per sampling time at 2, 3.5, 5, 7 and 9 hpt. Dashed curves represent the fit of the reduced model with logistic equation and solid curves show the fit of the full model (***Equation 1***– ***Equation 6***).

**Figure 4.**
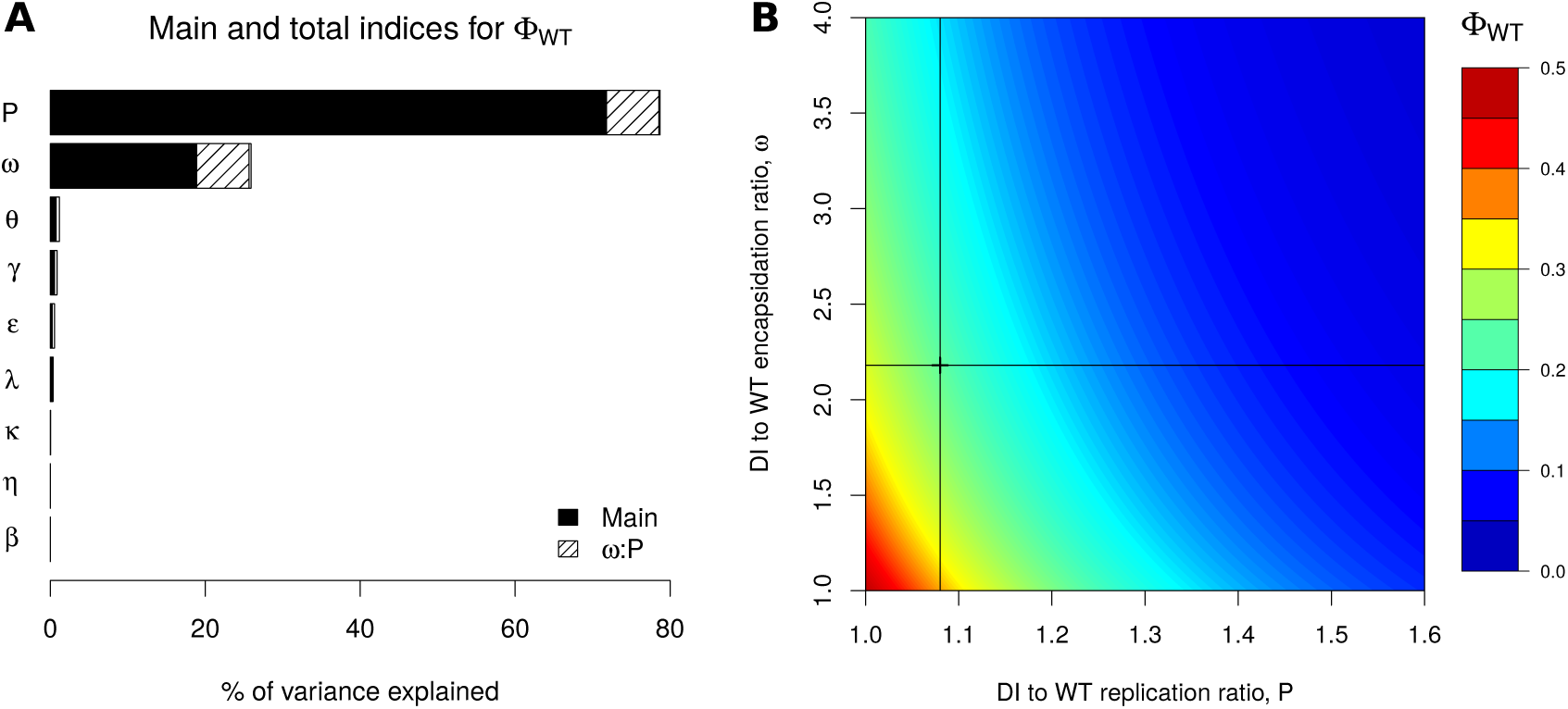
Sensitivity indices for the proportion of wild-type (WT) virions at cell lysis (Φ_*WT*_) and the impact of the most important parameters. A: Main, total and most important interaction indices for Φ_*WT*_. Main indices correspond to single parameter effect (black parts), and total indices add the effect of the factor in interaction with all other factors (second-order interactions, full bars). Hatched areas represent the importance of the strongest pairwise interaction between defective interfering (DI) to WT encapsidation rate *ω* and DI to WT replication factor *P* B: Heat map representing the impact of *P* and *ω* on Φ_*WT*_, all other parameters being fixed to their best estimated value. The little cross and lines in (B) show estimated parameter values from fitting the model to the experimental data.

## Model predictions

The aim of our work is to understand the competition dynamics of WT and DI genomes during co-infection. To achieve this goal, we used the model described above to study how changes in parameter values around their experimental estimates can impact the outcome of the competition. Additionally, we also used the model to evaluate the effect of initial infection conditions, such as initial genome copy numbers of WT and DI and a time delay of cell infection on their respective burst sizes.

### Sensitivity analysis

A sensitivity analysis was performed to identify parameters that have a significant effect on the output variable of interest, which we chose to be the proportion of WT and DI virions at the time of cell lysis, Φ_*WT*_ (***Equation 12***). Parameters were varied by ±50% of their best fit value based on experimental data, with five equally spaced values for each parameter (***Table 1***). The decay rate of genomes was not varied, because it was estimated to be negligible. The results indicate that the DI-to-WT replication ratio, *P,* and the DI-to-WT encapsidation ratio, *ω*, as well as their secondorder interaction, were the most influential factors for the variation of Φ_*WT*_, explaining 72%, 19% and 7% of the variance, respectively (***Figure 4***A). All the remaining factors and their second-order interactions had a negligible effect (less than 1% of the variance). Hence, our model predicts that only parameters associated with the DI design have a strong impact on the degree of suppression of WT by DI.

To further examine the effect of *P* and *ω*, we varied both parameters from their best estimated value (***Figure 4***B). As expected from the global sensitivity analysis, *P* was found to be more important than *ω* for the production of WT virions, the gradient of Φ_*WT*_ being steeper along *P*-axis than along *ω*-axis. Within the tested range of parameters *P* and *ω*, the value of Φ_*WT*_ varied between 2% and 50%. The reference value of Φ_*WT*_ corresponding to best-fit parameter estimates from experimental data was 23%. Therefore, we can predict that a DI particle with a lower replication factor, or, to a lesser extent, with a lower encapsidation rate than the DI particle used in the present work would weaken its competitivity with the WT virus, potentially leading to an increase in the proportion of WT virions at cell lysis of up to 27%. Conversely, a DI particle characterized by a higher replication factor or a higher encapsidation rate would strengthen its competitivity, potentially leading to a decrease in Φ_*WT*_ of up to 21%.

**Figure 5.**
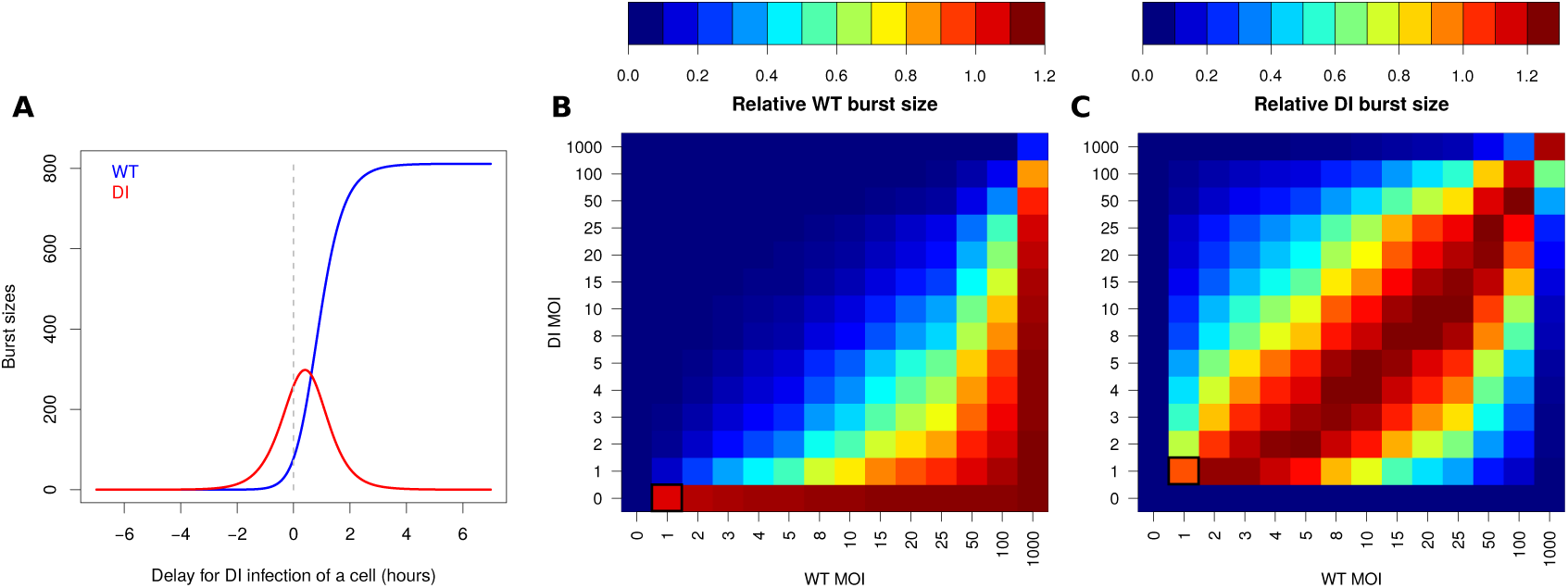
Impact of infection delay and initial multiplicity of infection (MOI) on wild-type (WT) and defective interfering (DI) burst sizes. A: WT (blue) and DI (red) burst sizes (encapsidated genomes at 9 hours post infection) for various delays in DI infection of a cell. Negative delays correspond to cases where DI infects first and positive delays to cases where WT infects first. The dashed gray line marks the case of simultaneous infection by WT and DI. B-C: Heat maps representing the relative burst sizes of (B) WT, ℬ_*WT*_, and (C) DI, ℬ_*DI*_, as a function of initial WT (x-axis) and DI (y-axis) MOIs (input). WT burst sizes were normalized by WT burst size resulting from WT:DI = 1:0 MOIs and DI burst sizes by DI burst size resulting from WT:DI = 1:1 MOIs (black squares).

### Effect of the multiplicity of infection and the timing of co-infection on WT burst size

We considered the impact of varying initial conditions including (i) the time difference between WT and DI infection of the cell and (ii) the initial quantities of WT and DI on the WT and DI burst sizes, ℬ_*WT*_ and ℬ_*DI*_, respectively. First, the time for DI infection compared to WT was varied from −7 to +7 hours post WT virus infection (***Figure 5***A). Negative delay values indicate that DI infects first, while positive values indicate that WT infects first. WT was allowed to produce at least one virion at cell lysis when DI was infecting the cell no more than 1.37 hours prior to the WT virus (delay = −1.37 hours). From this delay value, WT burst size increased as a steep logistic function, reaching a plateau at ℬ_*WT*_= 811 from a delay for DI infection of around 4 hours post WT infection. The curve of DI burst size was bell-shaped, reaching a maximum of ℬ_*DI*_ = 298 at a delay for DI infection of 0.4 hours post WT infection. The delay window allowing DI genome to be encapsidated and produce at least 1 virion is narrow, from −3.43 to 4.37 hours. Most importantly, DI burst size superseded WT burst size only until the delay of 0.62 hours. The difference between DI and WT burst sizes was the largest when DI infected the cell 0.03 hours prior to WT (ℬ_*DI*_ – ℬ_*WT*_= 181).

We then investigated the impact of varying initial WT and DI multiplicities of infection (MOIs, corresponding to the number of viral genomes successfully entering a cell and initiating infection) from 0 to 1000 on WT and DI relative burst sizes (***Figure 5***B-C). WT burst sizes were normalized by WT burst size obtained for WT:DI = 1:0 MOIs, while DI burst sizes were normalized by DI burst size obtained for WT:DI = 1:1 MOIs. WT relative burst size varied from 0 to 1.11 depending on MOIs. DI relative burst size ranged from 0 to 1.26.

At the optimal initial conditions maximizing WT burst size, WT must infect the cell with a larger MOI than DI. On the other hand, the optimal initial conditions for maximizing DI burst size are when WT is initially present in slightly larger quantity than DI. The DI needs enough WT to exploit its capsids and produce virions. Because (i) the DI replicates and encapsidates faster than the WT, (ii) only the WT produces free capsids and (ii) replication and capsid production result in resource depletion, it is more optimal for DI virion production to have the WT infect a cell in slightly higher quantity than the DI. In that case, the WT has a slight initial advantage over the DI and can use resources to produce free capsids. In return, the DI can exploit those free capsids at its own advantage as it replicates and encapsidates more efficiently.

We also examined the cross-effect of the time delay and the variation of initial MOIs (***Figure 5***– ***Figure Supplement 1***A). Globally, the shapes of WT and DI burst size curves as a function of delay are very similar between different WT and DI MOIs. The observed effect is a shift of the curves on the delay axis. Equal WT and DI MOIs generate very similar WT and DI burst size curves as a function of delay time. At these equal MOIs, DI competes more efficiently than WT upon co-infection, i.e. simultaneous infection of the cell by DI and WT (***Figure 5***–***Figure Supplement 1***A,B) and maximal DI burst size occurs when DI infects the cell around 0.3-0.5 hours after the WT (***Figure 5***–***Figure Supplement 1***A). As WT MOI gets larger than DI MOI, both the delay time that maximizes the DI to WT burst size difference and the peak of DI burst size shift towards delays where DI infects the cell before the WT (***Figure 5***–***Figure Supplement 1***A-C). On the opposite, as DI MOI gets larger than WT MOI, the shift is towards delays where WT infects the cell before the DI.

## Discussion

We used a combination of mathematical modeling and empirical measurements to get a better understanding of the mechanisms of interaction between a cooperator, the WT, producing capsid proteins as public goods, and a cheater, the DI, exploiting those capsid proteins from the WT when co-infecting a cell. In co-infected cells, WT and DI also compete for shared resources necessary for replication, a phenomenon known as exploitation competition (***Read and Taylor, 2001; Mideo, 2009; Bashey, 2015***). In a numerical analysis, we identified the critical parameters for the outcome of the intracellular competition in terms of the proportion of WT virions released at cell lysis.

***Huang and Baltimore*** (***1970***) argue that defective particles may influence the development and course of certain viral diseases. DI RNAs are produced through abnormal replication events and during high-multiplicity passage of the original virus (***Perrault, 1981***). Internal sequences of the original vRNA of DI RNA segments are deleted, whereas holding certain 5’ and 3’ end-specific sequences of the progenitor vRNA. A recent study on naturally occurring immunostimulatory defective viral genomes (iDVGs) reveals that they are generated during respiratory syncytial virus (RSV) replication and are strong inducers of the innate/natural antiviral immune response to RSV in mice and humans (***Sun et al., 2015***).

## Mechanism of defective interference

The precise mechanism of defective interference in poliovirus has been largely unclear, although there have been several classical studies evaluating DI particles (***Cole et al., 1971; Lundquist et al., 1979***). We specifically examined the competition between WT and DI genomes during RNA replication and the consequences of DI capsid exploitation for WT virus production (***Figure 1*** & ***Figure 4***).

Our experiment has provided three main results. First, the number of WT and DI genomes is lower in dually infected cells compared to singly infected cells (***Figure 1***C*i*& *ii*), indicating a limiting resource for replication. Second, in dually infected cells, DI genomes replicate faster than WT genomes (***Figure 1***C*iii*), showing the advantage of their shorter genome size (***Holland, 1991; Chao and Elena, 2017***). Third, the decrease in WT encapsidated genomes from singly to dually infected cells is two folds larger than that of WT naked genomes (***Figure 1***D*i*& C*i*), indicating that DI genomes, by trans-encapsidating in capsid proteins produced by the WT, further inhibit WT virions production.

In their experiment, ***Cole and Baltimore*** (***1973***) observed equivalent amounts of viral RNA produced in WT singly-infected cells or in dually-infected cells. In our experiment we obtain the same result, with 19,281 WT genomes in singly-infected cells at 9 hours post transfection vs. 18,153 WT and DI genomes in dually-infected cells (ratio of singly to dually-infected cells of 1.06).

We have designed a minimal mathematical model able to capture key features of the DI/WT interaction during a single-cell replication cycle. We accounted explicitly for depletion of cellular resources and available capsid proteins, the latter solely produced by the WT virus. This has allowed us to accurately describe the reference *in vitro* experimental data, and to predict new data on which the model had not been trained. In particular, the data fitting procedure has provided us with the possibility of estimating model parameters within biologically realistic ranges.

We expected the DI genome to replicate faster than the WT by a factor of approximately the ratio of WT to DI genome lengths, that is 7515bp/5733bp = 1.311. However, the best optimized value of the corresponding parameter *P* was lower (1.075, ***Table 1***). This discrepancy is most probably due to the time for various processes linked to replication to take place. For example, poliovirus replication first relies on the recruitement of membranes from intracellular organelles (endoplasmic reticulum, Golgi and lysosome) into clusters of vesicles, forming replication organelles (***den Boon and Ahlquist, 2010***). Then, replication starts by the synthesis of a negative strand of RNA, which serves as a template for positive-strand synthesis (***Novak and Kirkegaard, 1994***). The time for each of these preliminary steps to occur might lower the difference in replication speed between the WT and DI genomes. Additionally, the maximum number of replicases per RNA strand might be lower for the DI genome as it is shorter, lowering its overall replication speed compared to the WT (***Regoes et al., 2005***).

Experimental data suggests that some limiting resources are required and depleted during replication of viral genomes. Those resources might be phospholipids or lipids recruited by the viral machinery for the formation of replication complexes (***Altan-Bonnet, 2017; Nchoutmboube et al., 2013***) or host proteins involved in viral replication (***Regoes et al., 2005***). When both WT and DI co-infect a cell, they are in competition for the exploitation of those shared and limiting resources. As DI replicates faster (parameter *P* in the model), it depletes resources faster than WT, affecting WT replication compared to WT-only infections (***Bashey, 2015***). Additionally, ***Regoes et al.*** (***2005***) argued that virus genome replication represents a heavy burden for the cell, leading to its pathology and eventual death, and consequently to a slowdown in replication as the virus depletes resources. From fitting the models to the experimental data, we could estimate an initial replication rate of 3.02 − 3.07 • 10^−2^min^-1^, that can be assimilated to a case where the resources are not limiting. Then, as resources are depleted, the replication rate is predicted to decrease towards 0 logistically (***Figure 3***–***Figure Supplement 1***F).

## Insights on model fit to experimental data

The reduced model with the logistic equation underestimates the number of viral genome copies in singly infected cells. The logistic equation constrains the time-dependent genomic replication to be the same in both dually and singly infected cells. In other words, this reduced model assumes that depletion of resources required for replication is identical in singly or dually-infected cells. Hence viral genomes do not accumulate less when the other entity is co-infecting. As a result, genome copies are well predicted in dually-infected cells but underestimated in singly-infected cells. On the opposite, the full model is able to recapitulate the impact of resource depletion on RNA genome production in both dually and singly infected cells, as a specific variable for resources and mass action terms are added.

The full model overestimates the number of encapsidated WT genomes in singly infected cells. This result suggests that the WT virus encapsidates less efficiently when it is alone than when it is co-infecting a cell with the DI replicon. For the sake of simplicity, our model assumes that the encapsidation rate of WT is the same in singly- and dually-infected cells (*κ*), yielding the pointed-out overestimation. One hypothesis for this observation is that products of the DI could benefit to the WT when both co-infect a cell, enhancing WT encapsidation. Indeed, during co-infection, two viruses can exploit a common pool of resources equally (***Novella et al., 2004***). Our model predicts that the DI genomes encapsidate more efficiently than the WT, by a factor *ω* (= 2.185), and the factors enhancing DI encapsidation might also help WT encapsidation. In particular, the DI genome encodes non-structural proteases (3CβCD) which cleave P1 capsid protein precursor into several parts (***Burns et al., 1989; Ypma-Wong et al., 1988***), and additional expression of proteases by the DI may enhance WT capsid formation. Hence the assembly of WT capsids might be increased when the DI replicon is present in the cell and produces additional 3CβCD proteases. An additional hypothesis comes from the observation that replication and packaging of poliovirus are functionally coupled (***Nugent et al., 1999***). Hence the faster replication of DI might enhance encapsidation of the WT and trans-encapsidation of the DI.

A surprising observation is that the number of encapsidated genomes tend to decrease on average from 7 to 9 hours post transfection (from 410 to 385 for WT in singly infected cells, from 115 to 81 for WT in dually infected cells, and from 355 to 336 for DI in dually infected cells). This decrease is not recapitulated by the model as it does not include a decay term for encapsidated genomes. As the experiment was conducted at the cell population level and then the measurements were divided by the number of successfully transfected cells, it is possible that some cells got lysed before 9 hours post transfection, explaining a slowdown in virion production. However, we observe a decrease rather than a slowdown, which could be due to a decay of the virions, or to the infection of next cells by newly produced virions after cell lysis.

The cross-validation experiment showed an overall good predictive power of the model, although it underestimated the relative WT output when the DI was transfected in lower quantities than the WT virus (DI-to-WT input ratio of 0.46, ***Figure 3***–***Figure Supplement 2***). Full model simulations with best parameter values overestimates the number of WT encapsidated genomes in singly infected cells while the estimation is accurate in dually infected cells (***Figure 3***C & D). By normalizing the WT output by its value for WT only infection in the cross-validation experiment, we force the fit on the WT single infection condition, resulting in underestimations of WT relative output in dual infection condition. This can explain the observed discrepancy. It should be emphasized that, even with differing measurements between experiments and simulations (see Material and Methods), the model is able to recapitulate WT outputs for various inputs, which reinforces its robustness and predictive ability.

## Identification of parameters affecting WT virion production

A sensitivity analysis showed that the most important parameters for the proportion of WT virions at cell lysis when a cell is co-infected by the WT and the DI are DI relative replication factor (*P)* and encapsidation rate (*ω*) (***Figure 4***A-B). These are the only two parameters solely related to the DI construct. Modifications to the DI genome enhancing its replication and/or encapsidation could lead to a decrease in the proportion of WT virions at cell lysis from 23% to 2%. This result highlights the crucial role of exploitation competition (***Read and Taylor, 2001; Bashey, 2015; Mideo, 2009***) for resources necessary for genome replication and of interference competition (***Schoener, 1983***) for capsid proteins produced by the WT on the final proportion of WT virions at cell lysis.

## Impact of initial conditions

Overall, our model predicts a crucial impact of temporal spacing and order of infection, as well as initial MOIs, on the proportion of WT virions at cell lysis (***Figure 5*** and ***Figure 5***–***Figure Supplement 1***). At equal MOIs, the DI particle needs to infect a cell within approximately a 2 hours window before or after the WT in order to produce DI virions, and up to approximately 30 minutes after the WT in order to outcompete the WT in terms of burst sizes (***Figure 5***A). When simulaneously coinfecting a cell, the DI particles will maximise their virion production when WT and DI initial MOIs are approximately equivalent, and the WT particles will maximise their virion production when WT MOI is larger than DI MOI (***Figure 5***B-C). At equal MOIs, the difference between DI and WT virion production is maximised at approximately simultaneous co-infection of a cell. This difference maximisation is shifted towards cases where the DI infects a cell before the WT when WT MOI is larger than DI MOI. Conversely, it is shifted towards cases where the WT infects a cell before the DI when DI MOI is larger than WT MOI (***Figure 5***–***Figure Supplement 1***).

These results are in agreement with those of ***Cole and Baltimore*** (***1973***), who found that the extent of interference, assessed by the yield of WT poliovirus, is inversely proportional to the percentage of DI in the inoculum, and that it is also affected by varying the time interval between primary and secondary infection of a cell. In their viewpoint article, ***Read and Taylor*** (***2001***) highlight the importance of initial conditions, such as relative initial frequencies, temporal spacing and order of inoculation on the evolution of a population. An earlier review (***Henle, 1950***) also emphasized this aspect, while focusing on the exclusion of one virus strain by another. Experimental studies have shown inhibition of superinfection by a resident strain, in bacteria (***Berchieri and Barrow, 1990***) and in viruses (***Hart and Cloyd, 1990***). Another experiment on bacteria showed that the inhibition of superinfection was dose-dependent and also depended on the order of inoculation (***Lipsitch et al., 2000***). It is also known that picornaviruses rapidly induce resistance of the host cell to superinfection by the same virus, most probably because of inactivation or internalization of poliovirus receptors (***Koch and Koch, 1985***).

Importantly, ***Nugent et al.*** (***1999***) found that preaccumulated replicon RNAs are not trans-encapsidated by capsids made from a coinfecting helper virus, showing that only newly synthesized poliovirus RNAs are packaged. This could mean that, when the DI is the first to infect a cell, the genomes replicated before WT superinfection would not get trans-encapsidated by WT capsids. Hence our predictions might overestimate the DI burst size when DI is the first to infect a cell. This assumption would need to be tested in future experimental work.

### Limited resource and co-evolution

The competition between WT poliovirus and DI particles within cell can be analysed in light of evolutionary game theory. For a game between WT cooperators and DI defectors, the pay-off matrix features a fitness of zero in the case of a population composed only of DIs, because they are unable to reproduce (***Turner and Chao, 1999***). With such a feature, the evolution of a mixed WT and DI population is predicted to result in a polymorphic equilibrium, despite the greater pay-off that would result if the population was composed only of WT cooperators (***Turner and Chao, 1999***). These strategies of cooperation and defection are common in viruses, as co-infection of the same host cell induces competition for shared intracellular products (***Turner and Chao, 1999***). Evidence of such co-infections exists *in vivo*, as reported in the 2006 outbreak of dengue in India, where nearly 20% of infections comprised multiple dengue serotypes (***Bharaj et al., 2008; Mideo, 2009***). Long-term transmission of defective dengue viruses was also found in virus populations in humans and Aedes mosquitoes (***Aaskov et al., 2006***). Defective viruses were suggested to increase the overall incidence of transmission by modifying the virulence-transmissibility trade-off (***Ke et al., 2013***).

Resource availability can have important consequences on the dynamics and evolution of mixed pathogen populations (***Wale et al., 2017b; Turner and Chao, 1998; Read and Taylor, 2001***). For example, playing on resource availability could allow to slow the evolution of resistance to antimicrobial drugs (***Wale et al., 2017a***). In the case of DI particles, their presence within-cells infected by a WT virus is decreasing the number of resources available to the WT for replication and encapsidation, lowering WT virions production. Hence DI particles could be used to control WT infections, by lowering WT viral load, thereby facilitating further action of the immune system and/or drugs to clear the infection (***Huang and Baltimore, 1970***).

### Perspectives

We could take advantage of the features of DI particles to develop a new type of therapeutic antiviral strategy based on defective interference particle competition (***Frensing, 2015***). Our model suggests that parameters ***P***, the DI-to-WT replication ratio, and *ω*, the DI-to-WT encapsidation ratio, are the first and second most important parameters impacting the proportion of WT virions at cell lysis. Therefore, a rational strategy to strengthen interference activity of DI genomes and thus reduce the production of WT virions is to modify DI genomes towards higher replication speed and encapsidation efficiency. Such improvements may be realized by taking advantage of the evolvability of DI genomes. Serial co-passages of WT and DI particles followed by genetic analyses would allow for the screening of mutations providing higher replication or encapsidation of the DI. Also, the production of shorter DI genomes could lead to its faster replication.

Improving the interference at the intracellular level may cause less inhibition of WT viral load at the intercellular level, as there could be trade-offs. A reduced production of WT particles within-cells could result in a decreased MOI of WT viruses for the next infection cycle, and also to a decreased probability of a cell being co-infected. Furthermore, the narrow window of delay of co-infection for the DI to outcompete the WT as shown in ***Figure 5***A also suggests the importance of simultaneous infection. Interestingly, recent studies show several possibilities for how co-infection is or can be favored (***Robinson et al., 2014; Chen et al., 2015; Erickson et al., 2018***). Notably, the existence of vesicles containing multiple copies of virions as well as bacteria binding virions may increase the probability of simultaneous co-infection (***Robinson et al., 2014; Chen et al., 2015***). ***Erickson et al.*** (***2018***) reported that poliovirus binds lipopolysaccharide of bacteria, allowing co-infection of mammalian cells even at a low MOI.

While ecological studies for the control of pathogen populations mainly focus on preventing or slowing down the emergence of drug resistance (***Day et al., 2015; Roux et al., 2015; Wale et al., 2017a***) or on the evolution of virulence (***Frank, 1996; Brown et al., 2009***), we take an original approach here by rather studying how to use cheater defective pathogens, competing more efficiently for shared resources, for the control of disease-inducing pathogens. Since we learned the mechanisms of intracellular interference, in a future work we would like to apply these findings for the study of the competition between WT and DI at the larger level of the tissue, embedding intracellular knowledge. It would allow us to draw guidelines to optimize our DI construct at this level, based on WT viral load inhibition, and further confirm its efficiency *in vivo*.

## Material and Methods

### Competition experiment between defective genomes and wild-type genomes

#### Cells

HeLaS3 cells (ATCC CCL-2.2) provided by R. Geller and J. Frydman (Stanford University) were maintained in 50% Dulbecco’s modified Eagle medium and 50% F-12 medium (DMEM/F12) supplemented with 10% newborn calf serum (NCS), 100 U/ml penicillin, 100 U/ml streptomycin and 2 mM glutamine (Invitrogen).

#### Construction of viral cDNA plasmids

The cDNA plasmid prib(+)XpA, encoding the genome of poliovirus type 1 Mahoney strain under T7-promoter and hammerhead ribozyme sequences, was reported previously (***Herold and Andino, 2000***). Plasmid prib(+)XpA was digested by *NruI* and *SnaBI* (New England Biolabs) and ligated to produce prib(+)XpA lacking the poliovirus capsid-encoding region from 1175 to 2956 (prib(+)XpAΔ-1175-2956).

#### *In vitro* RNA transcription

Plasmids prib(+)XpA and prib(+)XpA-1175-2956 were digested by *EcoRI* or *PvuII*. Linearized plasmids were used as templates to obtain WT and DI genomic RNAs by *in vitro* transcription using RiboMAX™ Large Scale RNA Production Systems (Promega). *In vitro* transcribed RNAs were purified by phenol-chloroform extraction and the quality of purified RNAs was analyzed by electrophoresis on a 1% agarose gel in tris-acetate-EDTA Buffer (TAE).

#### Transfection of defective interfering and wild-type genomes

Monolayer of HeLaS3 cells was trypsinized and washed three times in D-PBS. Cells were resuspended in 1 ml D-PBS and the number of cells were counted on a hemacytometer, followed by adjusting the concentration to 1 × 10^7^ cells/ml. 800 *μ*l of cells and virus RNAs (5*μ*g of WT genomes and/or different amounts of DI genomes described later) were combined in a chilled 4-mm electroporation cuvette and incubated 20 minutes on ice. Cells were electroporated (voltage = 250 V, capacitance = 1000 *μ*F) using Gene Pulser II (Bio-Rad), washed two times, and recovered in 14 ml prewarmed (37 °C) DMEM/F12 medium with 10% NCS. Samples were distributed on 24 well plates (250 *μ*l/well).

Samples were collected at different time points (0, 3, 6, 9 hours for titration, and 0, 2, 3.5, 5, 7, 9 hours for RNA extraction) after electroporation. For titration and evaluation of encapsidated RNAs, samples were then frozen and thawed three times, followed by centrifugation at 2,500 g for 5 minutes, and supernatants were collected. Samples for evaluation of encapsidated RNAs were further treated with mixture of RNase A (20 *μ*g/ml) and RNase T1 (50 U/ml) (Thermo Fisher Scientific) for three hours. Samples were stored at −80 °C.

#### Titration of virus samples

Monolayers of HeLaS3 cells in 6-well plates were infected with 250 *μ*l of serially diluted virus samples at 37 °C for 1 hour and then overlaid with DMEM/F12 including 1% agarose. After 48 hours of infection, infected cells were fixed by 2% formaldehyde and stained by crystal violet solution. Titers were calculated by counting the number of plaques and multiplying their dilution rates.

#### RNA extraction

250 *μ*l of samples was added to 750 *μ*l of TRI-reagent LS (Sigma Aldrich), and RNAs were extracted following the kit protocol. Briefly, 200 *μ*l of chloroform was added to each sample, shaken vigorously, and incubated at room temperature for 10 minutes. Then samples were centrifuged at 12,000 g for 15 minutes at 4 °C. The upper aqueous phase was transferred to a fresh tube and 0.5 ml of isopropanol was added. After incubation at room temperature for 10 minutes, samples were centrifuged at 12,000 g for 8 minutes at 4 °C to precipitate RNAs at the bottom of the tube. The supernatant was removed and the residue was washed by 1 ml of 75% ethanol. After centrifugation at 7,500 g for 5 minutes at 4 °C, the RNA pellets were dried for 5-10 minutes. RNAs were resuspended in nuclease-free water.

#### Reverse transcription

2.5 *μ*l of RNA samples was mixed with 0.5 *μ*l of 2 *μ*M primer (5’-CTGGTCCTTCAGTGGTACTTTG-3’), 0.5 *μ*l of 10 mM dNTP mix, and 2.5 *μ*l of nuclease-free water. Samples were incubated at 65 °C for 5 minutes, and then placed on ice for 1 minute. After adding 10 *μ*l of cDNA synthesis mix (1 *μ*l of 10× RT buffer, 2 *μ*l of 25 mM MgCl2, 1 *μ*l of 0.1 M DTT, 1 *μ*l of RNaseOUT and 1 *μ*l of Superscript III RT enzyme), samples were incubated at 50 °C for 50 minutes, and then at 85 °C for 5 minutes to terminate reactions. 1 U of RNase H was added to each sample, followed by incubation for 20 minutes at 37 °C. Then, 0.1 U of Exonuclease I was added to each sample, followed by incubation at 37 °C for 30 minutes and at 80 °C for 15 minutes to terminate reactions. cDNA samples were stored at −20 °C.

#### Design of primers and Taqman probes

Primers and Taqman probes for droplet digital PCR assay were designed with PrimerQuest Tool (Integrated DNA Technologies). The primers and probe for WT genomes are 5’-CCACATACAGACGATCCCATAC-3’, 5’-CTGCCCAGTGTGTGTAGTAAT-3’, and 5’-6-FAM-TCTGCCTGTCACTCTCTCCAGCTT-3’-BHQ1. The primers and probe for DI genomes are 5’-GACAGCGAAGCCAATCCA-3’, 5’-CCATGTGTAGTCGTCCCATTT-3’, and 5’-HEX-ACGAAAGAG/ZEN/TCGGTACCACCAGGC-3’-IABkFQ.

#### Droplet digital PCR assay

2 *μ*l of serially diluted cDNA samples was mixed with 10 *μ*l of 2× ddPCR supermix for probes (BioRad), 1 *μ*l of 20× WT primers/probe, 1 *μ*l of 20× DI primers/probe, and 6 *μ*l of nuclease-free water. 20 *μ*l reaction mix of each sample was dispensed into the droplet generator cartridge, followed by droplet production with QX100 droplet generator (Bio-Rad). Then PCR was performed on a thermal cycler using the following parameters: 1 cycle of 10 minutes at 95 °C, 30 cycles of 30 sec at 94 °C and 1 minute at 60 °C, 1 cycle of 10 minutes at 98 °C, and held at 12 °C. Positive and negative droplets were detected by QX100 droplet reader (Bio-Rad). The data was analyzed with the QuantaSoft^™^ Software (Bio-Rad).

### Model reduction

As our mathematical model (***Equation 1***-***Equation 4***) presents a classical problem of parameter identifiability, we built a lower dimensional model to solve this problem by assuming that the decrease in resources due to viral uptake for replication follows a logistic decreasing function. This assumption was verified by analyzing the curves of *R*(*t*)*θε* as a function of time on a first set of “blind” optimizations (data not shown). Thus, we can recast the model using the following lower dimensional description:

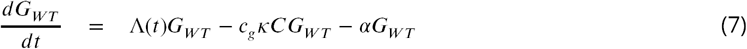

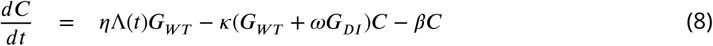

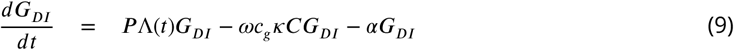

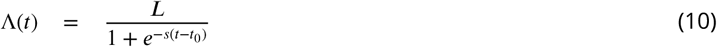

The logistic function (***Equation 10***) is characterized by the curve’s maximum value *L* and steepness *s*, and the time of the sigmoid’s midpoint *t*_0_. While this reduced version only decreases the number of parameters to be estimated by one (*L, s* and *t*_0_ instead of *θ, ε, λ*and *γ*), it partially solves the identifiability problem by removing biologically interpretable parameters and just assuming a logistic function for resource uptake and replication.

### Fit to experimental data

The model was fitted to the experimental data in order to estimate model parameters describing our biological system. Preliminary experimental data on the evolution of genome copy number showed that replication starts around 2 hours post transfection (data not shown). Indeed, there are several steps of poliovirus infection cycle before replication can start, including translation of positive-sense genomes (***Novak and Kirkegaard, 1994***) and transition from a linear, translating RNA to a circular RNA competent for replication (***Gamarnik and Andino, 1998***, ***2000; Herold and Andino, 2001; Schulte et al., 2015***). As our model does not account for those first steps, we only used experimental data from 2 hours post transfection for parameter estimation.

Raw experimental data are WT and DI total RNA copy number (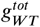 and 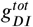) and RNAse treated genome copy number (*v*_*WT*_ and *v*_*DT*_). The former corresponds to the total number of genomes (naked and encapsidated) and the latter to the number of encapsidated genomes. The numbers of WT and DI naked (i.e. non-encapsdisated) genomes are thus: 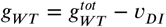 and 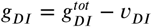. Additionally, as raw data was obtained at the cell population level, it was normalized by the average number of successfully transfected cells (data not shown) in order to get the average number of naked and encapsidated WT and DI genomes per cell.

In all, experimental data comprises three replicates of independent populations sampled at 2, 3.5, 5, 7 or 9 hours post transfection. Three different conditions were tested: (i) cells dually transfected by WT and DI genomes, (ii) cells transfected by WT genomes only and (iii) cells transfected by DI genomes only. Transfected volumes of WT and DI genomes were calibrated to a ratio of WT:DI = 4:1 in order to approximately obtain a ratio of 1:1 after transfection (preliminary experiment, data not shown).

Parameter estimation was achieved through nlminb optimization function in R software (***Gay, 1990***) embedded in an iterative process. Each optimization consisted in minimizing the sum of the least squares between experimental and simulated normalized data points for all variables and conditions. The least square function is as:

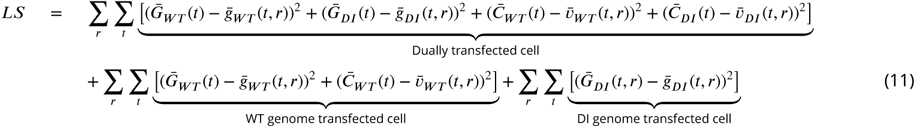

with *r* indicating the replicate number and *t* the sampling time (*t* = 3.5 to 9 hours post transfection). Initial conditions for all variables in all infection conditions were obtained from average experimental observations over the 3 replicates at *t*_0_ = 2 hours post transfection. In ***Equation 11***, we define:

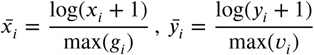

with *x*, resp. *y*, being either experimental (*g*, resp. *v*) or numerical (*G*, resp. *C*) naked, resp. encapsidated, genomes data and *i* for WT or DI.

The iterative process was applied as follows (see also ***Figure 3***–***Figure Supplement 3*** for a schematic representation). Boundaries on parameter values were defined based on a first set of “blind” optimizations, the intervals still remaining large and realistic. For each parameter *p*, let us denote *p*_*min*_ the lower boundary and *p*_*max*_ the upper boundary. For the first iteration, random values of parameters were drawn from uniform distributions, as *p*_*start*_ ∼ Unif(*p*_*min*_, *p*_*max*_), defining the starting point for optimisation. Let us denote 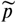 the optimised parameter value. In subsequent iterations, the starting point for each parameter was then randomly drawn from a uniform distribution, as 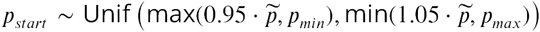. In all, 20 iterations were conducted, and this iterative procedure was implemented 250 times, each time with a different random starting point in the first iteration. Thus, 250 × 20 = 5000 optimizations were performed in total.

The goodness of fit was evaluated by ordinary least square (eq. ***Equation 11***) and sum of residuals *R*^2^ between experimental and simulated normalised data.

This optimisation procedure was applied in two steps. In the first step, the optimisation was performed on the reduced version of the model (***Equation 5***-***Equation 10***), thus estimating nine parameters: the six parameters corresponding to (i) the DI (*P* and *ω*), (ii) the production of capsids (*η*), (iii) the encapsidation process (*κ*) and (iv) the decay rates of genomes and capsids (*α* and *β*); and the three parameters of the logistic function representative of the time-dependent replication rate (*L, s* and *t*_0_). In the second step, the six redundant parameters between the reduced and full version of the model (*P, ω, η, κ, α* and *β*) were fixed to their best estimated value obtained during the first step. The remaining four parameters (*θ, ε, λ* and *γ*) were estimated by optimising the full version of the model (***Equation 1***-***Equation 6***).

### Model predictions

#### Cross-validation

We cross validate the results of our optimisation procedure by assessing how well the model is able to predict the relative WT virus burst size for various WT to DI initial ratios (after transfection) for which it has not been trained. We first obtain an optimal set of model parameters on our time series experimental data (*g* and *v*) featuring initial WT:DI = 1:1. We then test five additional DI-to-WT initial ratios, ranging from 0 to 3.6. Initial conditions for model simulations were set as the average of experimental values for each of the five initial ratios.

In the cross-validation experiment, evaluation of the relative WT virus burst size was based on the count of plaque-forming units (PFUs). In the time-series experiment that was used for parameter estimation, the number of WT virions (*v*_*WT*_) was estimated by digital droplet PCR. Assuming that the ratio of WT infectious to non-infectious particles and the multiplicity of infection (MOI) of WT virus are both constant independently of initial conditions, the relative PFU of WT virus for each initial condition should be a good proxy of the relative WT burst size.

The experiment was conducted on a cell population, and then the measurements were normalized by the number of successfully transfected cells. In some cases, the average experimental MOIs were small, potentially leading to not all cells being co-infected by WT and DI genomes. We integrated this aspect in our simulated burst size calculations, with the probabilities that a cell would be infected by both DI and WT genomes or only by WT genomes. We assume that the number of DI genomes infecting a cell *X*_*DI*_ results from a Poisson distribution of parameter the average DI MOI *n*_*DI*_, as *X*_*DI*_ ∼ *Pois*(*n*_*DI*_). The probability that no DI genome enters a cell is thus 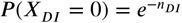. Conversely, the probability that at least one DI genome enters a cell is 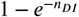. The expressions are equivalent for the WT virus. Let us denote the WT burst size in WT-DI infection as ℬ_*WT*_(*WT* -*DI*) and the WT burst size in WT-only infection as ℬ_*WT*_(*WT)*. We weighted WT simulated burst sizes as follows: 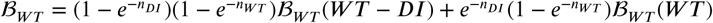.

WT PFU experimental values and WT burst size model predictions (ℬ_*WT*_) were normalized for all initial ratios by their respective values in the absence of DI genome (i.e. DI-to-WT initial ratio of 0). For the experimental data the average over the three replicates was taken for normalization. The performance of the model to predict relative WT burst size was evaluated by *R*^2^ and p-value of a Pearson correlation test between experimental and simulated datapoints.

#### Sensitivity analysis

A sensitivity analysis was performed to assess the relative importance of each parameter on the proportion of WT virions at cell lysis, defined as:

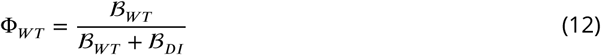

Based on parameter estimation, each parameter was approximately varied by ±50% of its best estimated value. Based on these boundaries, each parameter was allocated a vector of five equidistant values (except for *α* that was not varied because it was estimated at 0, see ***Table 1***). Then, all distinct combinations of parameter values were tested according to a full factorial design. In all, 5^9^ = 1, 953, 125 simulations of the full model were performed. All simulations started at time 0 hours post transfection with 10 copies of WT and DI genomes (*G*_*W T*_ (0) = *G*_*DI*_ (0) = 10), no capsids nor encapsidated genomes (*C*(0) = *C*_*W T*_ (0) = *C*_*DI*_ (0) = 0), and *R*(0) = *λ* /*γ*. Simulations were conducted until 9 hours post transfection. An analysis of variance (anova function in R software) was then conducted to assess the importance of each parameter and their second-order interactions on the variance of Φ_*WT*_.

#### Impact of delay and multiplicities of infection on WT and DI burst sizes

In the experiment, WT and DI genomes were co-transfected to cells and in quantities yielding an MOI ratio of approximately WT:DI = 1:1. We conducted two sets of simulations to study the impact of varying either (i) the time between cell infection by WT and DI or (ii) the MOIs of WT and DI on their burst sizes. All the simulations were conducted on the full version of the model (***Equation 1***-***Equation 6***). In the first set of simulations, WT and DI burst sizes were recorded for various delays between primary and secondary infection of a cell, ranging from −7 to +7 hours post transfection for the time of DI infection compared to the WT. The MOI upon infection of the cell was set to 10 for both WT and DI (i.e. *G*_*W T*_ (0) = 10 and *G*_*DI*_ (*t*_*d*_) = 10, with *t*_*d*_ the delay for DI infection), the number of capsids and encapsidated genomes to 0 and *R*(0) to *λ* /*γ*. In the second set of simulations, WT and DI burst sizes were recorded for various WT and DI initial MOIs, ranging from 0 to 1000. All the other variables were set as described for the study of delays. Then, WT burst sizes were normalized by the WT burst size corresponding to WT:DI = 1:0 initial MOIs (i.e. infection by the WT virus only at low MOI), and DI burst sizes by the DI burst size corresponding to WT:DI = 1:1 initial MOIs (i.e. infection by both WT and DIs at low MOI).

## Acknowledgments

This work is supported by the DARPA Intercept Program.

---

**Figure 3–Figure supplement 1.**
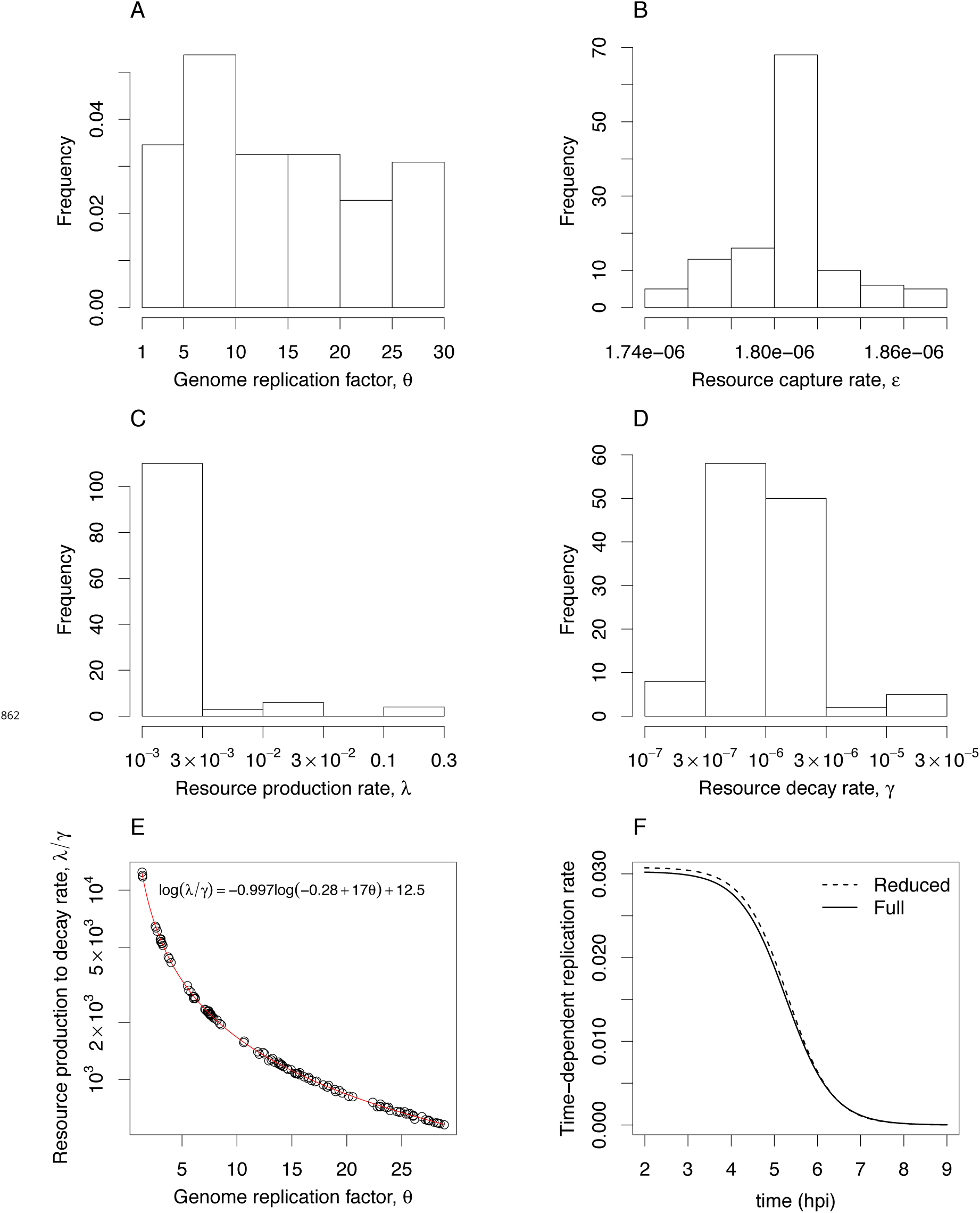
Histograms and correlations of the genome replication factor (*θ*), the resource linear production (*λ*), the decay rate (*γ*), and the resource capture rate (*ε*).

**Figure 3–Figure supplement 2.**
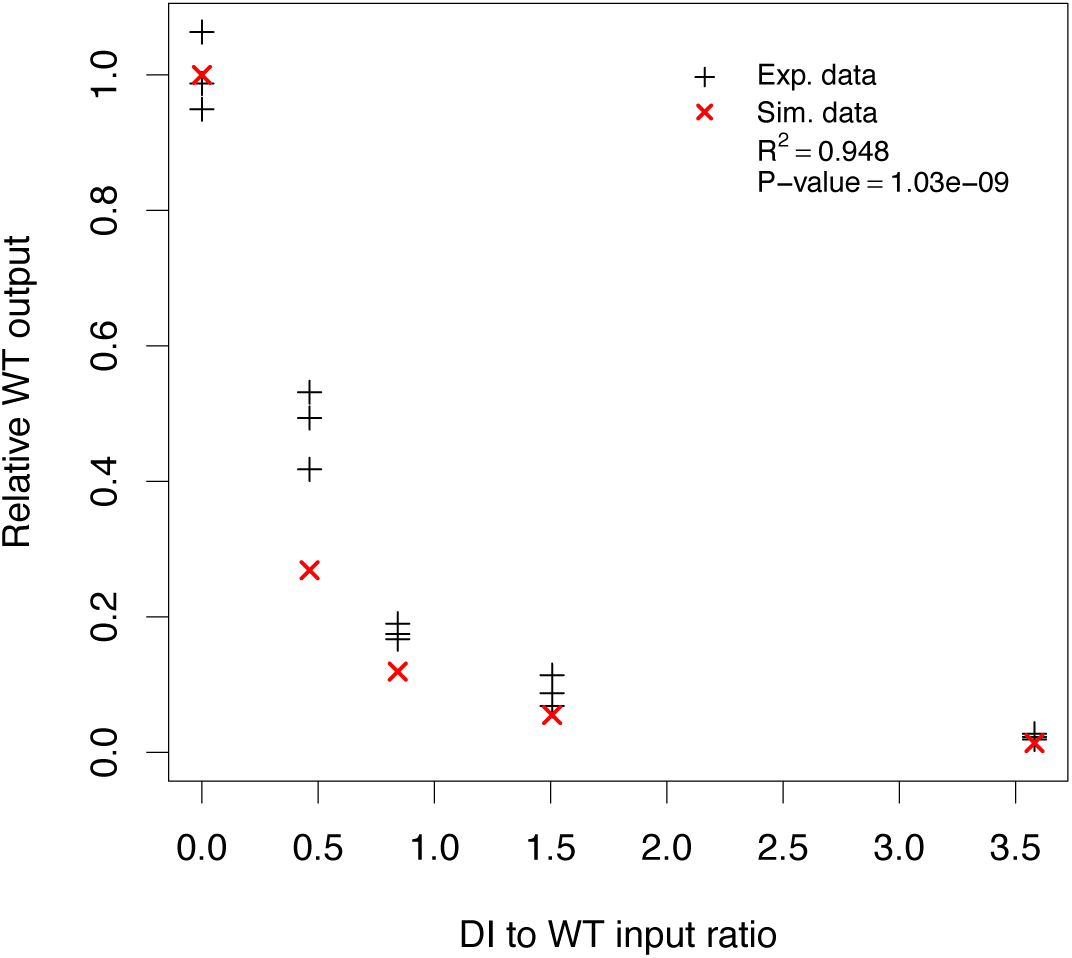
Cross-validation of the model (***Equation 1***– ***Equation 6***).

**Figure 3–Figure supplement 3.**
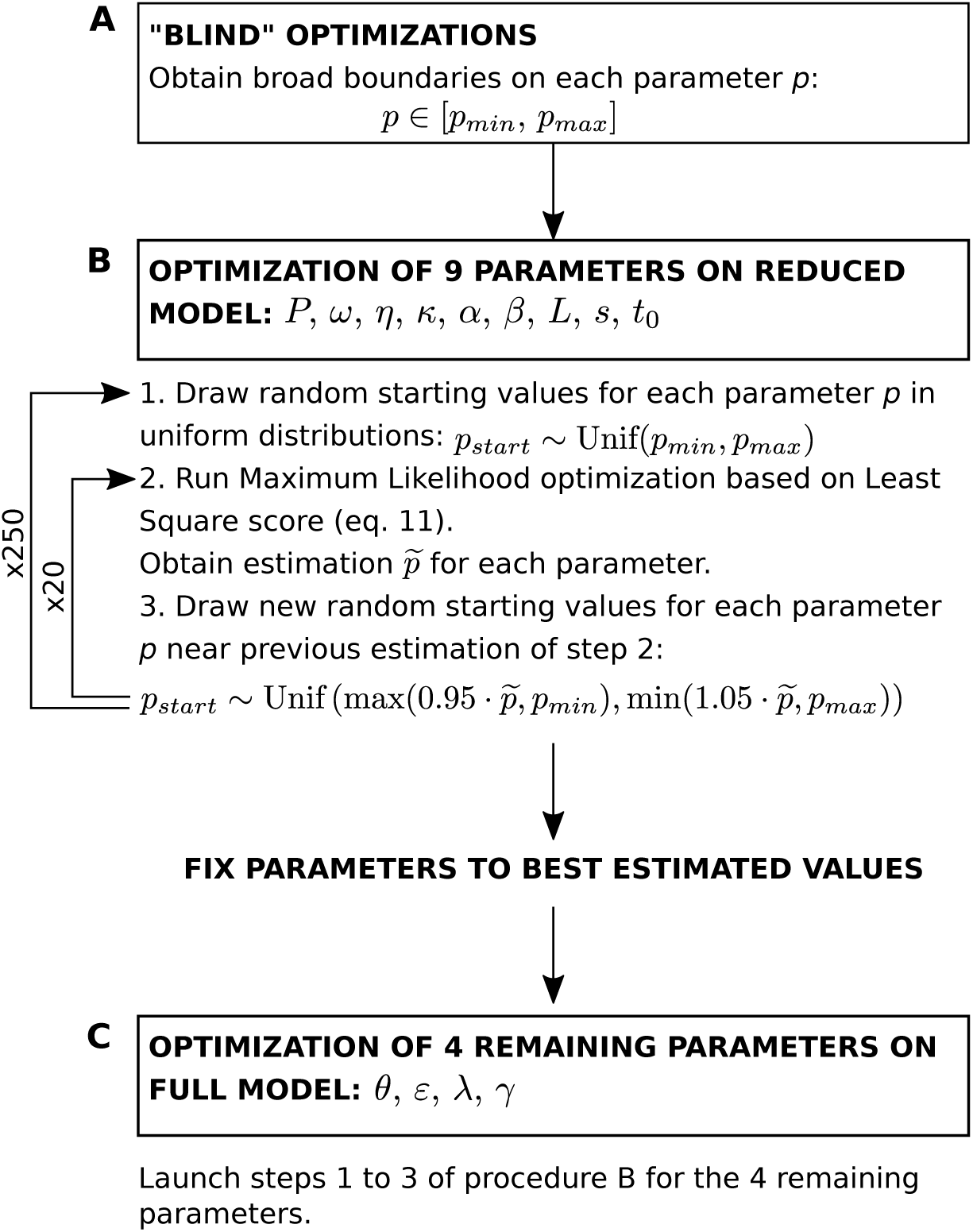
Diagram of the parameter fitting procedure.

**Figure 5–Figure supplement 1.**
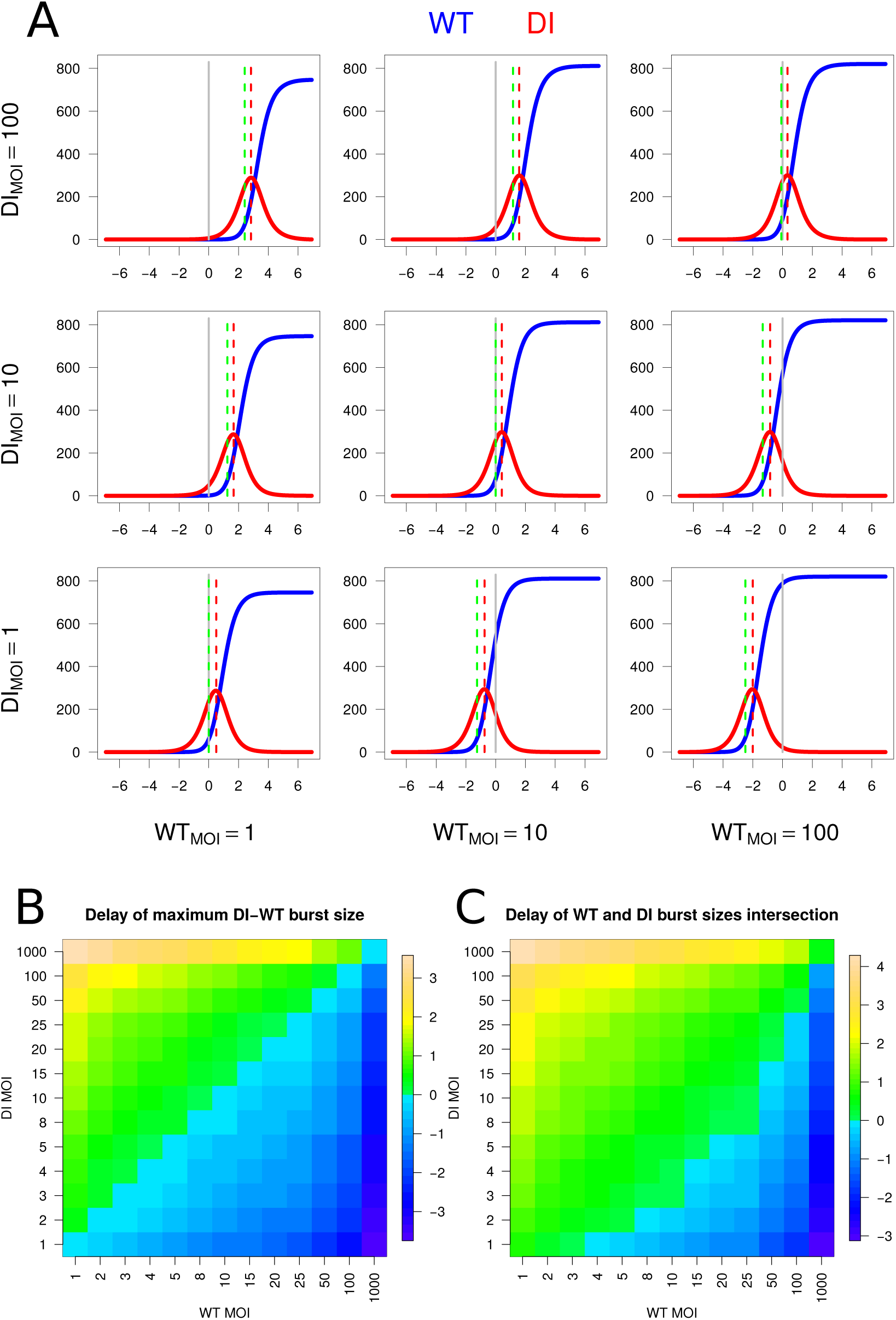
Delay at various multiplicities of infection (MOIs). A: Impact of delay for defective interfering (DI) particle infection of a cell (x-axis, in hours) on wild-type (WT, blue) and DI (red) burst sizes (y-axis). One line represents one DI MOI and one column one WT MOI. The grey vertical line indicates no-delay (simultaneous infection). The red vertical line indicates the peak of DI burst size and the green vertical line the maximum difference of DI to WT burst size. B: Heat map of the delay for the maximum difference of DI to WT burst size (green lines in A). C: Heat map of the delay for WT and DI burst size curves intersection.

